# Functional and evolutionary integration of a fungal gene with a bacterial operon

**DOI:** 10.1101/2023.11.21.568075

**Authors:** Liang Sun, Kyle T. David, John F. Wolters, Steven D. Karlen, Carla Gonçalves, Dana A. Opulente, Abigail Leavitt LaBella, Marizeth Groenewald, Xiaofan Zhou, Xing-Xing Shen, Antonis Rokas, Chris Todd Hittinger

## Abstract

Siderophores are crucial for iron-scavenging in microorganisms. While many yeasts can uptake siderophores produced by other organisms, they are typically unable to synthesize siderophores themselves. In contrast, *Wickerhamiella*/*Starmerella* (W/S) clade yeasts gained the capacity to make the siderophore enterobactin following the remarkable horizontal acquisition of a bacterial operon enabling enterobactin synthesis. Yet, how these yeasts absorb the iron bound by enterobactin remains unresolved. Here, we demonstrate that Enb1 is the key enterobactin importer in the W/S-clade species *Starmerella bombicola*. Through phylogenomic analyses, we show that *ENB1* is present in all W/S clade yeast species that retained the *<u>ent</u>*erobactin biosynthetic genes. Conversely, it is absent in species that lost the *ent* genes, except for *Starmerella stellata*, making this species the only cheater in the W/S clade that can utilize enterobactin without producing it. Through phylogenetic analyses, we infer that *ENB1* is a fungal gene that likely existed in the W/S clade prior to the acquisition of the *ent* genes and subsequently experienced multiple gene losses and duplications. Through phylogenetic topology tests, we show that *ENB1* likely underwent horizontal gene transfer from an ancient W/S clade yeast to the order Saccharomycetales, which includes the model yeast *Saccharomyces cerevisiae*, followed by extensive secondary losses. Taken together, these results suggest that the fungal *ENB1* and bacterial *ent* genes were cooperatively integrated into a functional unit within the W/S clade that enabled adaptation to iron-limited environments. This integrated fungal-bacterial circuit and its dynamic evolution determines the extant distribution of yeast enterobactin producers and cheaters.

## Introduction

Most organisms on earth require iron as a redox mediator that is key to numerous biological processes, including respiration; protection from reactive oxygen species; photosynthesis; and the biosynthesis of amino acids, lipids, deoxyribonucleotides, and sterols. Although iron is abundant in the crust of Earth, its availability to living organisms is generally low. This poor bioavailability occurs because iron exists mainly in its oxidized ferric (Fe^3+^) state, which is largely insoluble in water in neutral and basic pH environments. To circumvent iron shortage, microorganisms have evolved the capacity to scavenge iron from environmental stocks by secreting and taking up siderophores, a chemically diverse group of secondary metabolites that bind to Fe^3+^ with very high affinity and specificity (Haas et al. 2008; Kramer et al. 2020). These siderophore-mediated iron acquisition systems require active synthesis and secretion of siderophores in iron-free form, as well as specific transporter mechanisms to recognize and selectively take up the Fe^3+^-siderophore complex.

In contrast to most bacteria and filamentous fungi, yeasts of the subphylum Saccharomycotina (hereafter, yeasts) are generally unable to synthesize siderophores with a few exceptions, such as *Kluyveromyces lactis*, which produces pulcherrimin (Krause et al. 2018). The model yeast *Saccharomyces cerevisiae*, however, can utilize iron bound to various structurally distinct siderophores produced by other microbial species through two distinct systems (Yun et al. 2001). One system depends on the *FRE* gene family, which encodes plasma membrane reductases (Fre1, Fre2, or Fre3) that reduce and dissociate iron from siderophores at the cell surface; iron is then translocated through the plasma membrane by the high-affinity ferrous iron (Fe^2+^) transporter complex (Ftr1 and Fet3) (Stearman et al. 1996). Another system depends on four siderophore transporters of the major facilitator superfamily (MFS) that are distinct in substrate specificity and named Arn1, Arn2, Sit1 (also known as Arn3), and Enb1 (also known as Arn4) (Lesuisse et al. 1998; Heymann et al. 1999; Heymann et al. 2000a; Heymann et al. 2000b). This system allows a Fe^3+^-siderophore complex to enter the cells prior to any reduction step with high affinity and high specificity. Among these siderophore transporters, Enb1 is highly specific to the siderophore enterobactin and has previously only been reported in yeasts belonging to the genus *Saccharomyces*, whereas Sit1 transporters, which recognize a wide variety of siderophores, are abundant across the yeast phylogeny (Dias and Sá-Correia 2013). From an ecological and evolutionary perspective, siderophores can be shared between cells as public goods, and their production typically represents a form of cooperation (Julou et al. 2013; Weigert and Kümmerli 2017). However, yeasts with siderophore transporters can act as cheaters and exploit the benefits of siderophore production without paying the cost of synthesis and, thus, may outcompete the producers to a certain extent (Griffin et al. 2004; Dumas and Kümmerli 2012). The potential fitness benefits of being a cheater may have driven the evolution of siderophore transporter diversity in yeasts. While the biochemical properties of siderophore transporters have been examined by a wealth of studies in *S. cerevisiae* and other model yeasts (Yun et al. 2000; Ardon et al. 2001; Heymann et al. 2002), their evolutionary origins are largely unknown.

The exponentially rising amount of genomic sequence data have made yeast comparative genomics a powerful tool to explore the genetic bases and evolutionary histories of important traits (Butler et al. 2004; Hittinger and Carroll 2007; Marcet-Houben and Gabaldón 2015; Vakirlis et al. 2016; Gonçalves and Gonçalves 2019; Krause and Hittinger 2022). One of the most striking evolutionary events brought to light by comparative genomics is the horizontal operon transfer (HOT) of an *<u>ent</u>*erobactin biosynthesis pathway from an ancient bacterium in the order Enterobacteriales into the *Wickerhamiella*/*Starmerella* (W/S) clade of yeasts (Kominek et al. 2019). All the *ent* genes required for enterobactin biosynthesis were maintained and functionally expressed in at least some of the yeast species examined, which constitute a newly discovered and surprising group of enterobactin producers. However, genes encoding bacterial ATP binding cassette (ABC) transporters for enterobactin uptake were not found in these yeasts. The capacity to lock iron away from competitors would be counterproductive if the producer species were unable to utilize the iron bound to enterobactin, which suggests the presence of one or more alternative transporter mechanisms in the W/S clade yeasts.

Siderophore-mediated iron uptake is crucial for fungal iron homeostasis and virulence, but it has rarely been studied in yeasts other than *S. cerevisiae* and the opportunistic pathogen *Candida albicans* (Haas et al. 2008). Here, we investigate iron transport mechanisms using the enterobactin-producing yeast *Starmerella bombicola* as a model for targeted gene disruption and show that this species uses the Enb1 transporter for the uptake of enterobactin-bound iron. Through phylogenomic analyses, we show that *ENB1* is a fungal gene that was likely present in the most recent common ancestor (MRCA) of the Dikarya, the fungal clade that contains the phyla Basidiomycota and Ascomycota. This fungal gene was then functionally integrated with the horizontally acquired bacterial *ent* operon and evolved to operate coordinately in the W/S-clade yeasts as an adaptation to varying iron availability. Finally, we infer that a horizontal transfer of the *ENB1* gene from an ancestor of the W/S clade (order Dipodascles) to an ancestor of the order Saccharomycetales (which includes *S. cerevisiae*), along with multiple independent gene duplications and losses, shaped the patchy phylogenetic distribution of enterobactin cheaters in yeasts. This study highlights the intriguing interplay between fungal and bacterial genes, which formed a functional unit in some yeasts to produce a valuable resource, and later separated through dynamic evolution as cheaters arose to exploit that resource.

## Results and Discussion

### Enb1 imports enterobactin in *Starmerella bombicola*

While the HOT event and the subsequent genetic adaptations enabling bacterial gene expression and enterobactin synthesis in the W/S-clade yeasts are well-documented (Kominek et al. 2019), the mechanism by which these yeasts uptake the iron bound by enterobactin remains unknown. In the model bacterium *Escherichia coli*, the Fe^3+^-enterobactin complex is first translocated into the periplasm by the outer-membrane siderophore receptor FepA; it is then transported into the cytoplasm via the ATP binding cassette (ABC) transporter encoded by three genes: *fepC*, *fepD*, and *fepG*, which are parts of the *E. coli ent* operon (Usher et al. 2001; Raymond et al. 2003). However, neither *fepA*, *fepC*, *fepD*, nor *fepG* was found in the genomes of any W/S-clade yeasts, suggesting the presence of one or more alternative transport mechanisms.

We examined the possible mechanisms by which W/S-clade yeasts utilize enterobactin-bound iron using *St. bombicola* as a genetically tractable model species. Functionally related genes are often located in genomic proximity, even in eukaryotes (Hurst et al. 2004). The intercalation of genes encoding a eukaryotic ferric reductase (Fre) and an uncharacterized transmembrane protein (Tm) between two *ent* genes in a subset of the W/S-clade yeasts suggested a possible role in reductive uptake for this *FRE-TM* gene pair (Figure 1A). We also identified a homolog of the *S. cerevisiae ENB1* (*ScENB1*) in *St. bombicola* with 52% amino acid sequence identity and 96% coverage, which suggested the possibility that its function might be conserved as well. To determine which, if any, of these candidate genes were involved in the utilization of enterobactin-bound iron, we engineered two *St. bombicola* mutants by deleting either the *ScENB1* homolog (*enb1Δ*) or the *FRE-TM* (*fre-tmΔ*) gene cluster and compared their growth to the wildtype (WT) strain in both regular and low-iron media. The *fre-tmΔ* mutant exhibited comparable growth patterns to the WT in both media (Figure 1B), suggesting the *FRE-TM* gene pair is not essential for iron acquisition or could be functionally redundant with other ferric reductase genes, including homologs of *FRE1-FRE8* (Yun et al. 2001). Although it grew normally with regular iron concentrations, the *enb1*Δ strain showed minimal growth in low-iron conditions (Figure 1B). This result implies that the Enb1 homolog plays a crucial role in iron uptake in iron-limited conditions.

**Figure 1.**
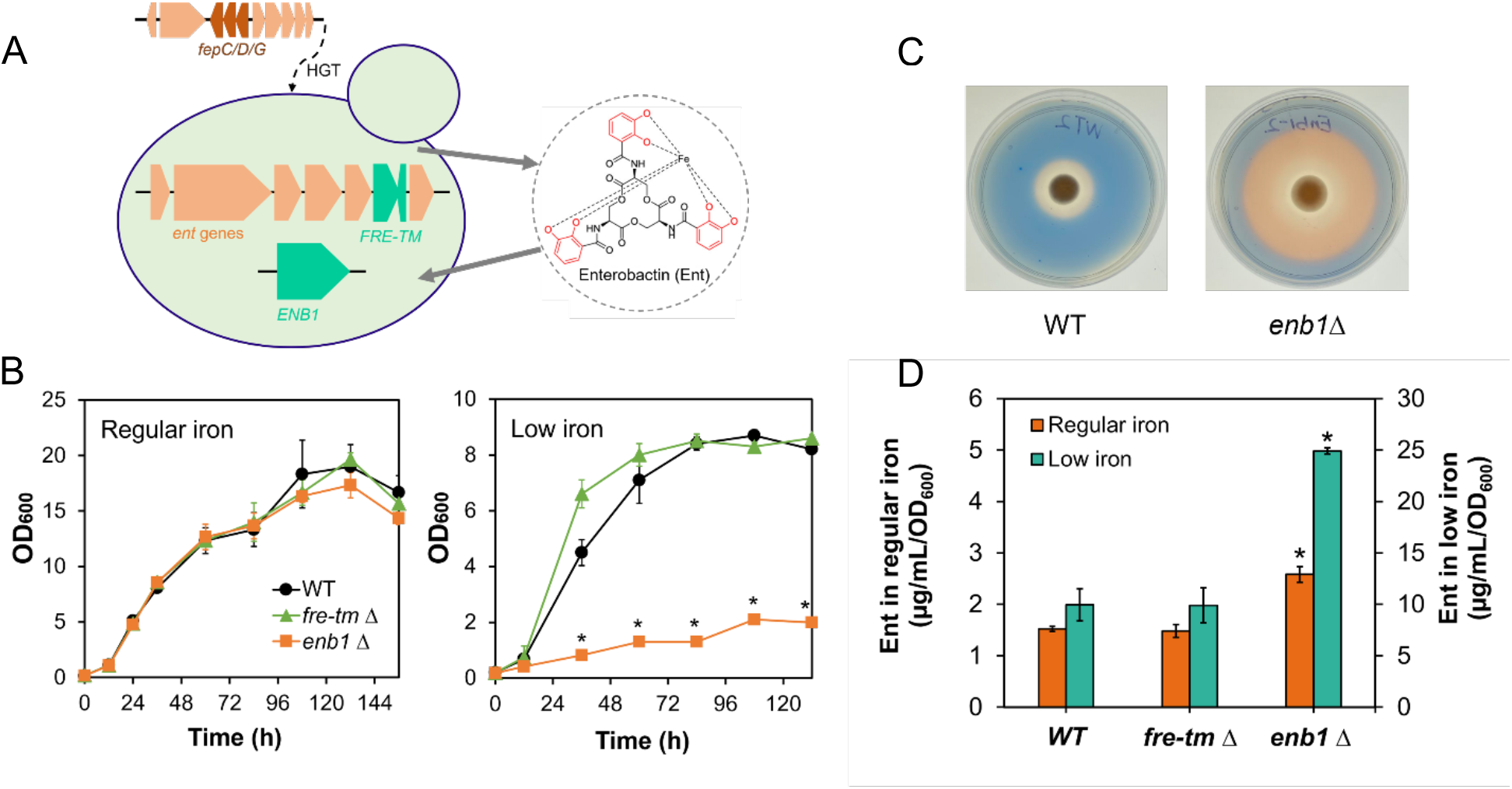
Enb1 is the key transporter mediating the uptake of enterobactin-bound iron in *St. bombicola*. Genes related to enterobactin biosynthesis and candidate genes for transport are native fungal genes (green) or underwent horizontal gene transfer (HGT) from a bacterial operon (salmon) (A). Growth of *St. bombicola* wildtype (WT) and mutants in regular- and low-iron conditions (B). The minimal growth of the *enb1ι1* strain in low-iron medium could be attributed to intracellular iron storage in vacuoles and other compartments during preculturing (Raguzzi et al. 1988) or reductive transport of enterobactin-bound iron with low affinity (Lesuisse et al. 2001). Chromeazurol S overlay (O-CAS) assay of the WT and *enb1ι1* mutants; iron sequestration converts the blue color to orange (C). Extracellular enterobactin accumulation of *St. bombicola* mutants (D). Data are presented as mean values and standard deviations of three independent biological replicates. Statistical significance of the differences between the *enb1ι1*, *fre-tmι1*, and WT strains was evaluated by Student’s *t*-Test. *, *p*-value < 0.001.

To delve into the function of the Enb1 homolog, we measured the levels of extracellular enterobactin. We found that all three strains produced substantially higher amounts of enterobactin in low-iron medium compared to regular medium (6.5-9.6-fold increase) (Figure 1D). This result indicates that enterobactin biosynthesis in *St. bombicola* increases under conditions of iron scarcity and might have evolved as an integral part of the regulatory network governing iron hemostasis. Moreover, the *fre-tmΔ*strain amassed similar levels of enterobactin as the WT in both culturing conditions, whereas cells of the *enb1Δ* strain accumulated 1.7-fold and 2.5-fold more enterobactin than the WT in regular medium and low-iron medium, respectively. An orthogonal colorimetric assay also showed a notably stronger enterobactin signal from the *enb1Δ*strain as compared to the WT strain (Figure 1C), which suggests that the mutant experiences severe iron starvation in low-iron conditions because it cannot effectively import enterobactin. Thus, we conclude that the *ENB1* homolog encodes a functional Enb1 transporter that is required for normal uptake of enterobactin-bound iron in *St. bombicola*.

Interestingly, disruption of neither *ENB1* nor *FRE-TM* noticeably affected the export of enterobactin. In *E. coli*, *entS*, as part of the *ent* operon, encodes the primary enterobactin exporter, which, together with the outer membrane channel TolC, mediates the secretion of enterobactin (Furrer et al. 2002). Since no homolog of *entS* could be identified in *St. bombicola* and other W/S-clade yeasts (Kominek et al. 2019), how the enterobactin synthesized in the cells is secreted to the medium remains an unanswered question.

### The origin of *ENB1*

To determine whether *ENB1* was horizontally acquired by the W/S-clade yeasts from bacteria like the *ent* genes, we performed BLASTp searches against the NCBI non-redundant sequences (nr) database using the amino acid sequence of *St. bombicola* Enb1 (*Stb*Enb1) as the query. Proteins from the Saccharomycetales, including *Sc*Enb1, were retrieved with *E* values equal to or lower than 5e-132, above which point proteins from filamentous fungi started to be recovered (Dataset S1-1). The *E* values obtained from some filamentous fungi hits were as low as 2e-117, suggesting strong sequence similarity with the *Stb*Enb1. Among the top 2,626 BLASTp hits recovered using an *E* value cutoff of e-60, only a few were from archaea or bacteria, while all the others were fungal proteins (Dataset S1-2). According to the maximum likelihood (ML) phylogeny constructed with these top hits (Figure S1), *Stb*Enb1 and its close homologs from the Saccharomycetales clustered monophyletically on a long branch with 100% bootstrap support, whereas the proteins from archaea and bacteria scattered across the larger gene tree on distant branches. Hence, we conclude that closely related Enb1 homologs are only found in fungi, and the *StbENB1* gene is unlikely to have been acquired horizontally from bacteria.

The high sequence similarity of *Stb*Enb1 to proteins from filamentous fungi inspired us to examine the evolutionary history of Enb1 in the kingdom Fungi. For this purpose, we conducted a HMMER scan for Enb1 homologs against a dataset containing 1,644 published fungal genomes (Li et al. 2021) using the aligned sequences of *Sc*Enb1 and *Stb*Enb1 as queries; we obtained 1,061 protein sequences by employing a bit-score cutoff of 300 and a *E*-value cutoff of 0.05 (Dataset S2). The resulting proteins were exclusively from the Dikarya; the vast majority of them were recovered from the subphyla Saccharomycotina (491) and Pezizomycotina (553), while only 17 of them were from the subphyla Agaricomycotina, Wallemiomycotina, and Taphrinomycotina. After accounting for putative gene duplication events that gave rise to distantly related paralogs (e.g. genes encoding Arn1, Arn2, and Sit1), the ML phylogenetic tree constructed using the 1,061 protein sequences (Figure S2) was generally consistent with the reported genome-scale species phylogeny (Li et al. 2021). Notably, *Sc*Enb1 and *Stb*Enb1, as well as Enb1 homologs from many yeasts belonging to the W/S clade and the order Saccharomycetales, formed a monophyletic group with 12 proteins from diverse fungi at 98% bootstrap confidence. The 12 closest relatives of yeast Enb1 were recovered from species that were scattered sparsely, but widely, in the Dikarya across the subphyla Pezizomycotina, Agaricomycotina, Wallemiomycotina, and Taphrinomycotina. Since this sparse clade roughly matched the species phylogeny, we propose that it comprises the extant Enb1 orthologs. Under this model, *ENB1* was likely present in the MRCA of the Dikarya, while its sparse and dispersed distribution is likely due to extensive losses in the various lineages.

Interestingly, the non-Saccharomycotina fungal species are either known plant pathogens, including *Eutypa lata*, *Rhizoctonia solani*, *Passalora fulva*, and *Ceratobasidium theobromae* (Thomma et al. 2005; Ajayi-Oyetunde and Bradley 2018; Ali et al. 2019; Lolas et al. 2020); or they can exhibit yeast-like growth, including three fission yeasts (*Schizosaccharomyces japonicus*, *Schizosaccharomyces octosporus*, and *Schizosaccharomyces cryophilus*), three dimorphic fungi (*Zymoseptoria brevis*, *Zymoseptoria ardabiliae*, and *Prillingera fragicola*) (Quaedvlieg et al. 2011; Takashima et al. 2019), and a basidiomycetous yeast (*Apiotrichum gamsii*) (James et al. 2016). This unusual phylogenetic distribution suggests a potential role of Enb1 in the evolution of fungal virulence and unicellular yeast growth, which is consistent with the previous finding that siderophore-mediated iron uptake is crucial for fungal iron homeostasis and virulence (Haas et al. 2008).

### Horizontal gene transfer of *ENB1* within yeasts

The presence of *ENB1* in *St. bombicola* and *S. cerevisiae*, which belong to two distantly related orders that diverged over 300 million years ago (Shen et al. 2018; Groenewald et al. 2023), prompted us to more thoroughly investigate its species distribution within the subphylum Saccharomycotina. We performed a HMMER search against 345 publicly available yeast genomes (Shen et al. 2018; Kominek et al. 2019) for Enb1 homologs using aligned sequences of Enb1 from *S. cerevisiae* (*Sc*Enb1) and *St. bombicola* (*Stb*Enb1) as the query. In addition to *Sc*Enb1 and *Stb*Enb1, 32 proteins, exclusively from the order Saccharomycetales and W/S clade of yeasts, were retrieved up to *E* value of 3.9e-170 and bit score of 565.5, after which point the *E* value ramped up and hit score dropped drastically (Dataset S3). All 34 closely related Enb1 homologs clustered monophyletically on a very long branch with 100% bootstrap support in the maximum likelihood (ML) phylogeny (Figure 2 & S3). Besides Enb1, we recovered the other three siderophore transporters, Arn1, Arn2, and Sit1, as well as the glutathione exchanger Gex1 and its paralog Gex2 (Dhaoui et al. 2011) from *S. cerevisiae*. Consistent with previous observations (Diffels et al. 2006; Dias and Sá-Correia 2013), *S. cerevisiae* Arn1, Arn2, Gex1, and Gex2, as well as all their homologs from other Saccharomycetales species, formed a single clade in the tree with 100% bootstrap support.

**Figure 2.**
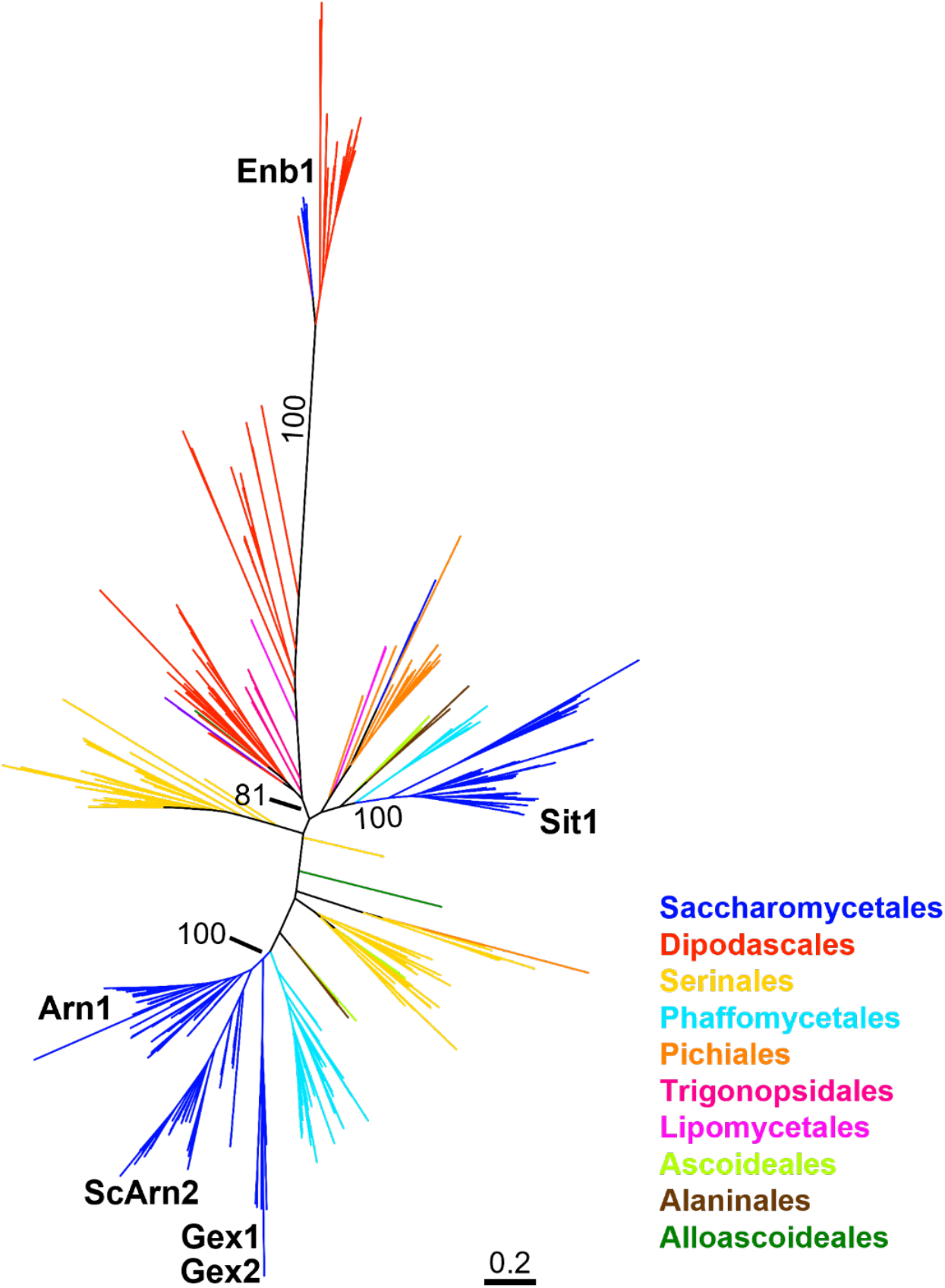
Maximum likelihood phylogeny of Enb1 homologs, including paralogs, from the yeast subphylum Saccharomycotina using an amino acid sequence alignment. Branch values shown are the percentages of branch support out of 1000 bootstrap replicates. The four Arn family siderophore transporters (Arn1, Arn2, Sit1, and Enb1) and two glutathione exchangers (Gex1 and Gex2) from *S. cerevisiae* are labeled in the tree. Yeast major clades are colored according to (Shen et al. 2018) and are now circumscribed as orders (Groenewald et al. 2023). Figure S3 shows a midpoint-rooted version of the same ML phylogeny; this rooting supports the conclusion that Enb1 is monophyletic.

**Figure 3.**
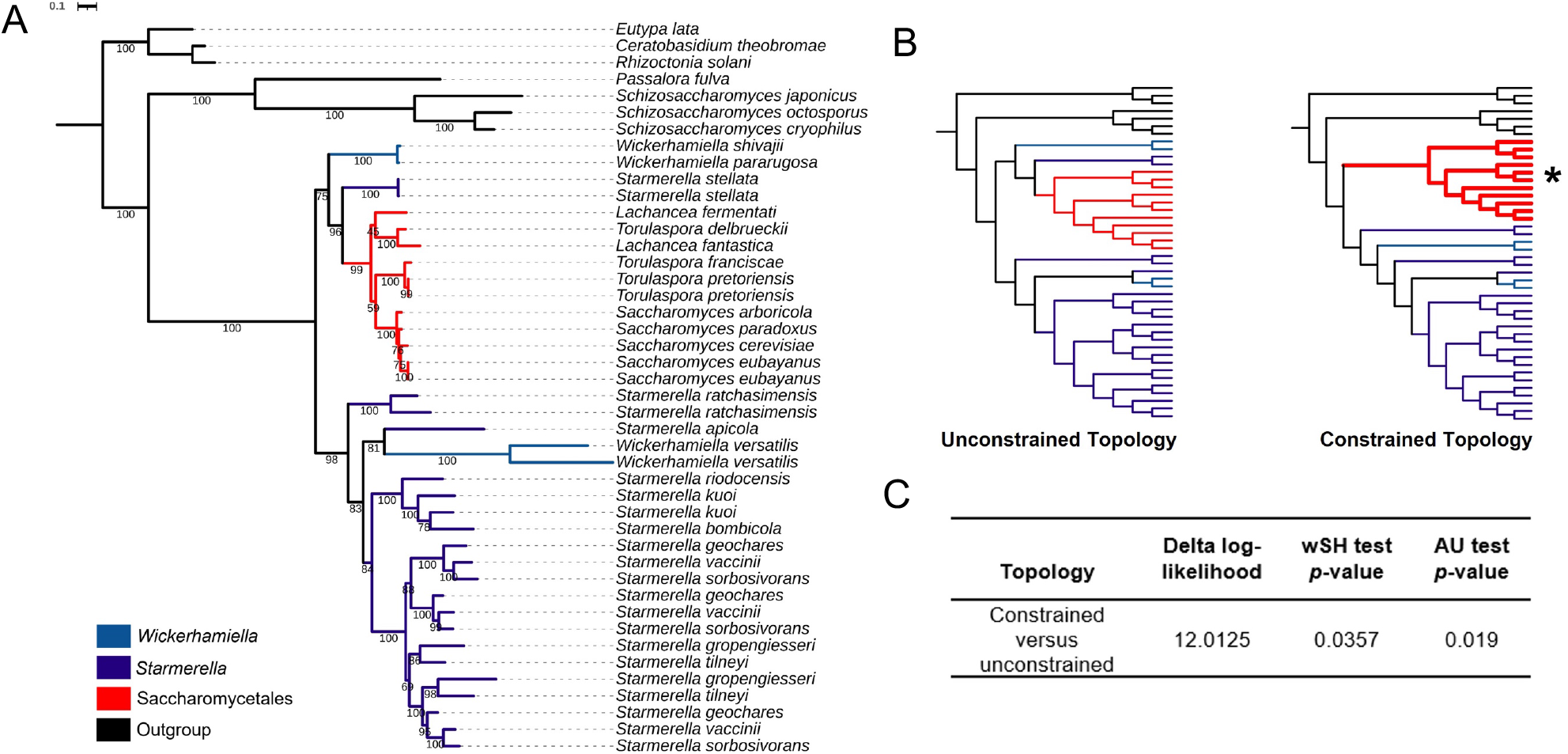
Evidence for HGT of *ENB1* in yeasts. Maximum likelihood phylogeny of the fungal Enb1 protein sequences most closely related to Saccharomycotina Enb1 (A), which are a clade from the full analyses shown in Figure S1. The 7 closest relatives of Enb1 proteins from filamentous fungi were chosen as outgroups. Branch values shown are the percentages of branch support out of 1000 bootstrap replicates. Branches are colored in light blue, dark blue, red, and black to represent species from the genera *Wickerhamiella*, *Starmerella*, order Saccharomycetales, and outgroup lineages, respectively. Branches with the same species name occur because some genomes encode multiple copies of Enb1. The simplified topologies of the unconstrained and constrained gene trees are shown in (B). Branches repositioned in the constrained topology are highlighted with wider lines and denoted with (*). The SH and AU topology test results are shown in (C). *p*-values of weighted Shimodaira-Hasegawa (wSH) test and approximately unbiased (AU) test were both lower than the statistical significance threshold of 0.05.

From these searches, we discovered clear *ENB1* orthologs in species from the genera *Saccharomyces*, *Torulaspora*, and *Lachancea*. Surprisingly, these *ENB1* orthologs from the order Saccharomycetales were nested within those of the W/S clade (Figure 2), which suggests the possibility of a horizontal gene transfer (HGT) event from the W/S clade in the order Dipodascales to the order Saccharomycetales. This model is consistent with the fact that previously known *ENB1* homologs were identified exclusively in the sub-telomeric regions of yeasts belonging to the genus *Saccharomyces* (Dias and Sá-Correia 2013), which constitute a favorable chromosomal location for the acquisition of foreign genes (Kellis et al. 2003; Novo et al. 2009). To better illuminate the evolution of Enb1 within yeasts, we identified three more Enb1 homologs from *Wickerhamiella shivajii* and *Starmerella stellata*, whose genomes were recently published (Opulente et al. 2023). Using maximum likelihood, we carefully examined the phylogenetic relationships between the 37 yeast Enb1-like protein sequences, as well as the 7 closest relatives of Enb1 from fungal outgroups. As expected, the yeast Enb1-like transporters formed a monophyletic clade with 100% bootstrap support (Figure 3A, Figure S2). However, there were at least 3 instances where the Enb1 tree did not recover the expected phylogenetic relationships between yeast taxa (Figure 3A &4). Most notably, Enb1 homologs from the Saccharomycetales were nested within the W/S clade and indeed were sister to the distantly related species *St. stellata* with 96% bootstrap support. This observation suggests the acquisition via HGT of an *ENB1* gene by an ancient Saccharomycetales from an ancestor of W/S clade yeasts. To test this hypothesis, we performed topology tests where we compared the likelihood of the ML-supported (unconstrained) topology against the likelihood of an alternative constrained topology that enforced the reciprocal monophyly of the Saccharomycetales- and W/S-clade homologs (Fig. 3B). Both weighted Shimodaira–Hasegawa (wSH) and approximately unbiased (AU) tests significantly rejected the null model of vertical inheritance and instead supported HGT (Fig. 3C).

**Figure 4.**
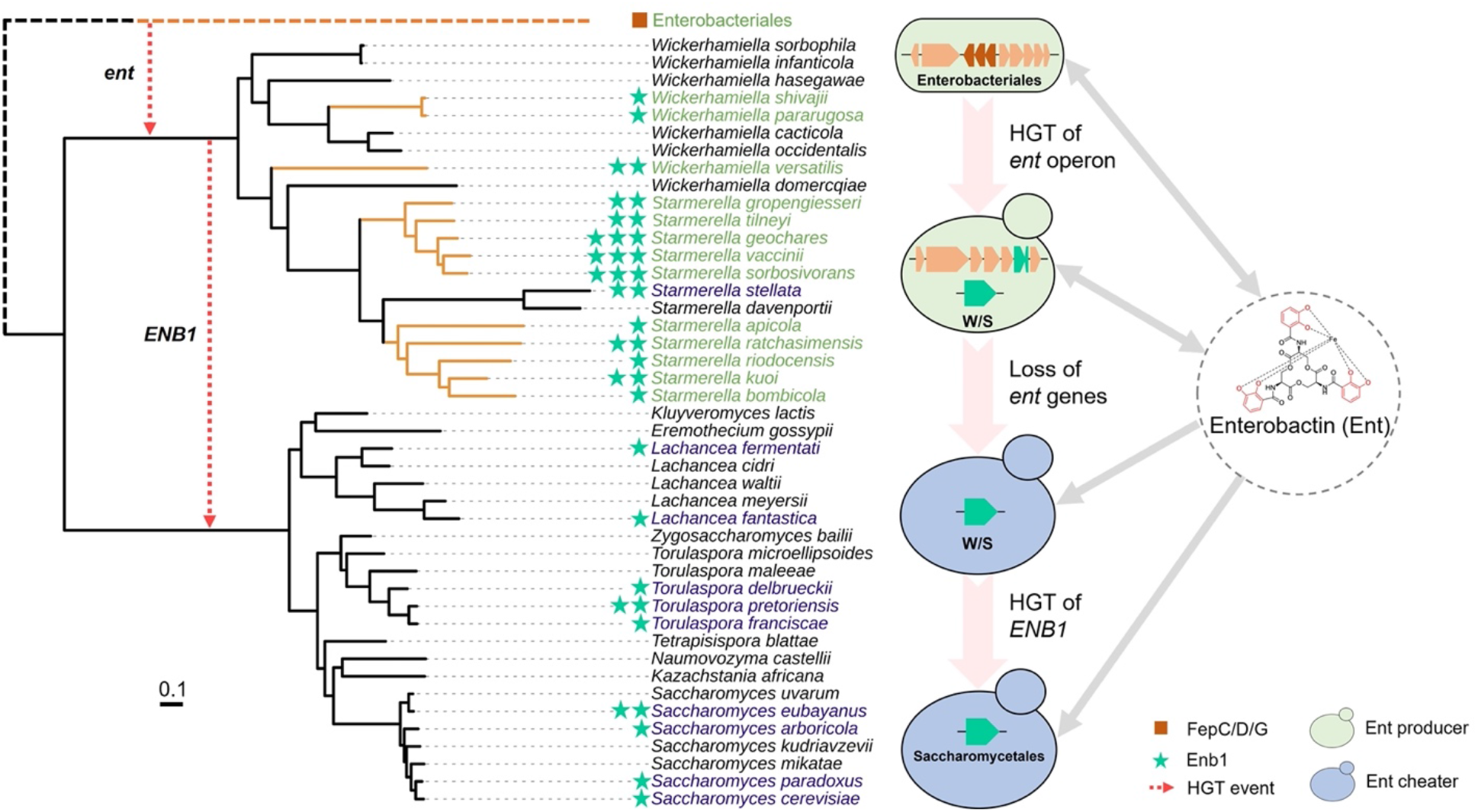
Evolution of the fungal enterobactin transporter encoded by *ENB1* and its functional integration with the bacterial *ent* operon in yeasts. The genome-scale species phylogeny was adopted from a recent publication (Opulente et al. 2023) by pruning irrelevant branches of the phylogeny of 1,154 yeasts using iTOL v5 (Letunic and Bork 2021). Branches colored in orange represent the presence of *ent* genes. Dashed arrows in red denote horizontal gene transfer (HGT) events. The brown square represents genes encoding the Enterobacteriales FepC/D/G ABC-type enterobactin transporter. Green stars represent the number of genes encoding fungal Enb1 transporters. Branch names in green represent enterobactin producers harboring both *ENB1* and *ent* genes. Branch names in blue represent enterobactin cheaters harboring only genes encoding the Enb1 transporter. Species with names in black have neither *ENB1* nor *ent* genes. Gene icons are colored as in Figure 1.

Second, the *St. stellata* Enb1 homolog did not associate as expected with other *Starmerella* branches of the tree in the Enb1 phylogeny (Figure S4A). Instead, *St. stellata* Enb1 formed a monophyletic clade with the Saccharomycetales, *Wickerhamiella pararugosa*, and *W. shivajii* homologs. This topology suggests another possible HGT event from this *Wickerhamiella* lineage into an ancestor of *St. stellata*. To test for additional HGT events between *Wickerhamiella* and *Starmerella*, we performed a series of topology tests. First, we enforced the placement of *St. stellata* Enb1 either as an outgroup of the *Starmerella* Enb1 clade (Constrained Topology 2, Figure S4B) or to its exact location in the species phylogeny (Constrained Topology 3, Figure S4B). The wSH and AU tests of Constrained Topology 3 robustly rejected the null model of vertical inheritance, while the same tests of Constrained Topology 2 did not reject the null model (Figure S4C). Since Constrained Topology 3 incorporates additional incongruences between the gene and species phylogenies, we conclude that HGT from an ancient *Wickerhamiella* into an ancestor of *St. stellata* is one of many evolutionary models that are consistent with the data, including more complex scenarios involving multiple HGTs, cryptic gene duplication, or incomplete lineage sorting within the W/S clade. Additionally, the topology tests did not significantly support a possible HGT associated with the *Wickerhamiella versatillis* Enb1 (Constrained Topology 1, Figure S4B, Figure S4C). Given the evolutionary distances involved, we conclude that *ENB1* has experienced a complex history in some W/S-clade lineages, including the lineage leading to *St. stellata*. We further conclude that one of these ancient W/S lineages likely acted as the donor of the Saccharomycetales *ENB1*.

### Duplications and losses of *ENB1* and its functional integration with a bacterial operon

To get more insight into the evolutionary mechanisms leading to the distribution of *ENB1* orthologs in yeasts, we mapped its presence, absence, and copy numbers onto a genome-scale species phylogeny inferred in a recent publication (Opulente et al. 2023) (Figure 4). Given its broad distribution in the clade, *ENB1* was almost certainly present in the MRCA of the W/S clade. It could have been acquired from other fungi by HGT, or *ENB1* could have been present in the MRCA of Saccharomycotina and then lost in all other known yeast lineages (at least 9 independent losses by maximum parsimony). The *St. bombicola enb1Δ* mutant phenotype (Figure 1) indicates that expression of an Enb1 transporter is essential to for normal uptake of iron bound to enterobactin. Thus, *ENB1* would convey a selective advantage in iron-limited environments where enterobactin is present, including enterobactin made by other microbes, but especially for organisms making their own enterobactin. Therefore, we hypothesize that the *ENB1* gene was actively expressed in the MRCA of the W/S-clade yeasts prior to the HOT event that introduced the *ent* operon into yeasts. Under this scenario, the *fepC/D/G* genes from the operon, which would have encoded the bacterial ABC-type enterobactin transporters, could have been lost easily due to functional redundancy with fungal *ENB1*.

The phylogenetic distribution of *ENB1* copy numbers and inference by maximum parsimony suggest at least two duplication events in the W/S clade: one that occurred before the divergence of *Wickerhamiella versatilis*, *Wickerhamiella domercqiae*, and the genus *Starmerella*; and a second that occurred in the last common ancestor of *Starmerella geochares*, *Starmerella vaccinii*, and *Starmerella sorbosivorans*. This model is generally consistent with the Enb1 protein tree (Figure 3A). Additionally, maximum parsimony suggests multiple loss events occurred both prior to and following the duplications, leading to reduced copy numbers or *ENB1* absence in some W/S-clade yeasts. Disproportionate secretion and uptake of enterobactin could also result in its extracellular accumulation, potentially reaching levels detrimental to the cells that produce it. We hypothesize that altering the copy number of the *ENB1* gene through duplications and losses might be an evolutionary strategy taken by the W/S-clade yeasts to balance the expression levels of the biosynthetic enzymes and the transporter, while adapting to new environments with different iron availability.

Copy number variation aside, the presence and absence of *ENB1* almost perfectly mirrors that of the *ent* genes encoding enterobactin biosynthesis pathway. Specifically, *ENB1* is present in all the W/S-clade yeasts that retained the *ent* genes, while it is absent in nearly all species that lost the *ent* genes with *St. stellata* as the only exception. The correlation between the presence of the *ENB1* and *ent* genes is likely driven by the fact that the loss of *ENB1* would be deleterious to yeasts actively producing enterobactin because they would sequester iron that they could not import, a phenotype we observed in the *St. bombicola enb1Δ*mutant (Figure 1B). The sole exception is *St. stellata*, which is the only W/S yeast that harbors the *ENB1* gene in the absence of *ent* genes, a genetic configuration known to create public goods cheaters in other siderophore systems (Wang et al. 2015; Krause et al. 2018). Regardless of whether *St. stellata* reacquired *ENB1* from an ancient *Wickerhamiella* species via HGT or not (Figure S4), it likely represents a reversion to the ancestral cheater state prior to the HOT event, either through simple loss of the *ent* operon or also through loss and subsequent regain of *ENB1*.

## Conclusions

Siderophores can readily bind iron, even in an iron-limited environment, thereby both reserving iron for producers and depriving iron from species that lack the matching transporter for uptake (Niehus et al. 2017; Schiessl et al. 2017). However, it has been repeatedly shown that non-producers possess conditional competitive advantages over the producers because they act as cheaters and exploit foreign siderophores without the cost of synthesis (Griffin et al. 2004; Jiricny et al. 2010; Dumas and Kümmerli 2012). The fact that the MRCA of the W/S clade was likely converted from a cheater for enterobactin into an enterobactin producer through HOT suggests that there were ecological and evolutionary circumstances where being a producer had an advantage over being a cheater. The HOT was proposed to have occurred in insect guts, which are believed to foster intense competition for iron among bacteria, yeasts, and the host itself (Barber and Elde 2015; Kominek et al. 2019). Yeasts with the capacity to produce their own enterobactin, which has exceptionally high affinity for Fe^3+^ (Kf = 10^51^) (Carrano and Raymond 1979), may have been better positioned to outcompete nonproducers and species that produce different siderophores, especially in communities lacking enterobactin producers. Thus, the cooperative integration of the fungal *ENB1* gene and the bacterial *ent* genes as a functional unit for sequestering iron would have substantially contributed to the fitness of yeasts in highly competitive, iron-limited environments. After enterobactin-producing W/S-clade yeasts had begun to radiate, *ENB1* was horizontally transferred into an ancient lineage of Saccharomycetales (Figure 3), which allowed many additional yeasts, including *S. cerevisiae* (Petra Heymann et al. 2000), to exploit the enterobactin produced by others. Several secondary losses of *ENB1* then led to its patchy distribution in the order Saccharomycetales.

Siderophore uptake has rarely been studied in nonconventional yeasts, but this study suggests it is subject to surprisingly complex eco-evolutionary dynamics. Here, we identified Enb1 as the key transporter mediating the uptake of enterobactin-bound iron in *St. bombicola*, one of the extant W/S-clade yeasts that acquired a complete bacterial enterobactin biosynthesis pathway via an ancient HOT. Through phylogenomic analyses, we showed that the *ENB1* gene is of fungal origin and has a broad, but patchy, distribution in the Dikarya. Along with the horizontally acquired bacterial operon, this fungal gene formed a mosaic functional unit that operates coordinately in many W/S-clade yeasts to scavenge iron from the environment. We also propose how cheaters arose through the secondary loss of the *ent* operon and/or HGT of *ENB1*. Multiple gene duplication and loss events punctuate the *ENB1* phylogeny and further suggest a dynamic eco-evolutionary history of enterobactin cheaters in yeasts. This story shows how fungal and bacterial genes can be functionally integrated in the same species for the production of a resource, as well as how eco-evolutionary forces can then separate them as cheaters arise to exploit that resource.

## Materials and Methods

### Construction of *St. bombicola* deletion mutants

Genetic manipulations were performed in the *St. bombicola* PYCC 5882 strain obtained from the Portuguese Yeast Culture Collection. Strains used in this study are listed in Dataset S5. All the oligonucleotides used for construction of *St. bombicola* mutants are listed in Dataset S6. The hygromycin-resistance cassette with a *St. bombicola GPD* promoter and *S. cerevisiae CYC1* terminator was amplified from the previously constructed plasmid PJET-ADH1-HYG (Gonçalves et al. 2018). Genomic DNA of *St. bombicola* PYCC 5882 was isolated and purified using a modified phenol:chloroform extraction method as previously described (Hittinger et al. 2010). For disruption of the *ENB1* and *FRE-TM* genes, two sets of primers were used to amplify ∼1 kb upstream and ∼1 kb downstream of their coding sequences with 30 bp overlaps to the hygromycin-resistance cassette. The *ENB1* and *FRE-TM* deletion cassettes were obtained by assembling the upstream and downstream fragments of each gene with the hygromycin-resistance cassette using the NEBuilder^®^ HiFi DNA Assembly Master Mix (Catalog# E2621), followed by PCR amplifications and cleanup using the QIAquick^®^ PCR Purification Kit. Phusion^®^ High-Fidelity DNA Polymerase (Catalog# M0530) was used for all the PCR amplifications.

*St. bombicola* was transformed with each deletion cassette using a previously described electroporation protocol (Saerens et al. 2011). The transformants were screened on YPD (10 g/L yeast extract, 20 g/L peptone, 20 g/L dextrose, pH 7.2) plates supplemented with 650 µg/mL hygromycin B (US Biological) and verified using colony PCR and Sanger sequencing.

### Growth and iron-binding assays

*St. bombicola* strains were inoculated from glycerol stocks, which were stored at -80 °C, to YPD medium and precultured for 3 days at 30 °C in culture tubes. 2% dextrose was used as the carbon source for all cell cultivations. For low-iron cultivations, 1.7g/L Yeast Nitrogen Base without Amino Acids, Carbohydrates, Ammonium Sulfate, Ferric Chloride, or Cupric Sulfate

(US Biological) was used to make the low-iron synthetic complete (SC) medium (“low-iron medium” in brief) without direct addition of any iron source, although it may inadvertently acquire ambient iron during the medium preparation and cultivation processes. The low-iron medium also consisted of 5 g/L ammonium sulfate, 2 g/L Complete Dropout Mix (US Biological), and 200 nM cupric sulfate. The regular SC medium (“regular medium” in brief) consisted of 200 µg/L (1.2 µM) ferric chloride with all other components kept identical. YP medium contains about 25 µM of iron (Du et al. 2012), which is approximately 20 times of the iron concentration in the regular-iron SC medium. To avoid substantial iron carryover from the YPD precultures, cells were washed twice in deionized water and reinoculated at an initial optical density (OD_600_) of 0.2 in culture tubes with regular- and low-iron media for a second preculture. After 4 days, the cells were again washed and reinoculated at an OD_600_ of 0.2 into the corresponding media for the growth experiments, which were conducted in 250 mL shake flasks with 50 mL of regular- or low-iron medium at 30 °C.

The chromeazurol S overlay (O-CAS) solution was made with chromeazurol S, ferric chloride hexahydrate, PIPES (free acid), and agarose as described previously (Kominek et al. 2019). Cells of the *St. bombicola* wildtype and *enb1Δ* strains were precultured in 5 mL YPD medium for 3 days, harvested, and resuspended in 5 mL deionized water. 5 µL of the resulting cell suspension was spotted on the center of 60-mm diameter petri dishes consisting of low-iron SC agarose (1% w/v) and incubated at 30 °C. After 7 days, 6 mL of the O-CAS solution were overlaid onto low-iron SC plates. The plates were sealed with Parafilm and put in the dark at room temperature for another 7 days before pictures were taken on an LED light box.

### Enterobactin quantification

Supernatants of yeast cultures were harvested by centrifugation at the end of the growth experiments. The amount of enterobactin was quantified using a Shimadzu LCMS8040 by injecting 10 µL of the filtered supernatants onto a Phemonenex Kinetex XB-C18 column (P/N: 00G-4605-E0, 100Å, 250 × 4.6 mm, 5 µ particle size) held at 50 °C. The mobile phase was a binary gradient of A) water and B) acetonitrile, pumped at 0.7 mL/min. The gradient program was set according to %B using linear gradients between set time points, starting by holding at 5%B for 1 min; then ramping up to be 10%B at 5 min and 95%B at 6.5 min; then held at 95%B until 8 min, at which point it was returned to 5%B at 11 min and held at 5%B until the end of the program at 15 min. The column eluent flowed through a photo diode array detector scanning from 250–600 nm and into a 2-way switch valve to send it to waste or the mass spectrometer. The mobile phase was diverted to waste from time 0.1 – 3.9 min to aid in keeping the electrospray ionization (ESI) source clean by preventing the buffer salts present in the media from entering the mass spectrometer. The column eluent was sent to the ESI source from time 3.9 – 15 min.

The dual ion source was set to operate in ESI negative mode, with 2.5 L/min nitrogen nebulizing gas, 15 L/min nitrogen drying gas, desolvation line temperature set to 250 °C, and the heating block temperature set to 400 °C. The mass spectrometer was programed to collect Q3-scans negative ions from 200 – 1000 *m/z* and, for the quantification of enterobactin, three multiple-reaction monitoring (MRM) transitions were acquired: 668→178, CE:53, relative intensity: 100; 668→222, CE:33, relative intensity: 53; and 668→445, CE:21, relative intensity: 22. Argon gas at 230 kPa was used for the MRM fragmentations. The retention time (10.2 min), relative intensities of the three MRM transitions, and a 7-point calibration curve from 125–10 µg/mL were determined using an enterobactin standard (Sigma Aldrich #E3910). To minimize carryover the needle was washed for 10 sec before and after each injection. Two blank methanol injections were made between each sample to ensure any carryover enterobactin was below the threshold of detection (∼7 µg/mL). As the cell growth varied substantially for the 3 strains grown in different conditions, the concentrations of enterobactin were normalized by the final cell optical density of yeast cultures for fair comparisons.

### Screening for the presence of *ENB1* homologs

Basic Local Alignment Search Tool protein (BLASTp) analyses in a publicly available genomic assembly of the *St. bombicola* PYCC 5882 strain (GeneBank ID: GCA_003033785.1) (Gonçalves et al. 2018) were performed using the Enb1 protein sequences from *Saccharomyces cerevisiae* (*Sc*Enb1) as the query. The identified *ENB1* ortholog in *St. bombicola* was functionally validated using targeted gene replacement as described above and designated as *StbENB1*. To ascertain whether Enb1-like transporters could be found outside the fungal kingdom, the *Sc*Enb1 amino acid sequence was used as BLASTp query against the NCBI non-redundant protein sequence (nr) database (Datasets S1-1 & S1-2). The hits with *E*-values < 2e-60 were filtered by removing redundant sequences with > 90% similarity using the clustering function of the MMseq2 (Steinegger and Söding 2017). The resulting 1165 sequences were used to construct a maximum likelihood (ML) phylogeny. We visualized and plotted phylogenetic trees using iTOL v5 (Letunic and Bork 2021). To evaluate the prevalence of *ENB1* homologs among fungal genomes, a hidden Markov model (HMMER) sequence similarity search (v3.3, http://hmmer.org) was conducted against all the protein-coding sequences pulled out from 1644 published fungal genome assemblies (Li et al. 2021) using the aligned sequences of *Sc*Enb1 and *Stb*Enb1 as the query. HMMER hits with *E*-values < 0.05 are listed in Dataset S3, and protein hits with bit-scores > 300 were retrieved for phylogenetic analyses.

We also identified homologs of Enb1 across the 345 published yeast genome assemblies (Shen et al. 2018)(Kominek et al. 2019) using a separate HMMER search. Protein annotations used for the HMMER search were generated previously by the MAKER genome annotation pipeline v2.31.8 (Holt and Yandell 2011; Shen et al. 2018; Kominek et al. 2019), again using the alignment of *Sc*Enb1 and *Stb*Enb1 sequences as the query. HMMER hits with E-values < 0.05 were listed in Dataset S3, and the 435 hits with bit scores > 300 were retrieved for constructing a maximum likelihood (ML) phylogeny (Figure 2). We similarly retrieved Enb1 homologs from the recently published genomes of *W. shivajii* and *St. stellata* (Opulente et al. 2023).

### Phylogenetic analyses and topology tests

Protein sequences of hits from the aforementioned HMMER and BLASTp analyses were retrieved to construct maximum likelihood trees to infer the phylogenetic relationships of *ENB1* homologs. MAFFT (Katoh and Standley 2013) v7.508 was used to align the sequence with the strategy L-INS-i. Initial inferences of the phylogenies were performed with IQ-TREE (Nguyen et al. 2015) v2.2.0.3 using ModelFinder (Kalyaanamoorthy et al. 2017) with automated model selection based on 1000 bootstrap pseudoreplicates. The LG+R10, LG+G4, LG+R7, and LG+R10 amino-acid substitution models were selected because they had the lowest Bayesian Information Criterion scores to construct the trees in Figure 2, Figure 3A, Figure S1, and Figure S2, respectively. The seven closest relatives of the Enb1 orthologs in filamentous fungi were chosen as outgroups for the Enb1 phylogeny (Figure 3). To investigate the likelihood of HGT events between the W/S clade and order Saccharomycetales, phylogenetic analyses of Enb1 proteins were repeated with specific constraints, and significance was assessed with AU and SH topology tests. The constrained tree in Figure 3B enforced the reciprocal monophyly of the Saccharomycetales and the W/S clade, while the constrained trees in Figure S4 enforced the repositioning of *St. stellata* and *W. versatilis*. The substitution model selected for all the constrained trees was LG+G4.

## Supplementary Materials

Supplementary data are available at *Molecular Biology and Evolution* online.

## Acknowledgments

We thank David J. Krause, Trey K. Sato, and members of the Hittinger and Rokas groups for helpful discussions. This material is based upon work supported in part by the Great Lakes Bioenergy Research Center, U.S. Department of Energy, Office of Science, Office of Biological and Environmental Research under Award Number DE-SC0018409; the National Science Foundation (under grant Nos. DBI-1906759 to K.T.D., DEB-2110403 to C.T.H., and DEB-2110404 to A.R.); and the National Institute of Food and Agriculture, United States Department of Agriculture, Hatch project 7005101. C.T.H. is an H. I. Romnes Faculty Fellow, supported by the Vice Chancellor for Research and Graduate Education with funding from the Wisconsin Alumni Research Foundation. Research in the Rokas lab is also supported by the National Institutes of Health/National Institute of Allergy and Infectious Diseases (R01 AI153356), and the Burroughs Welcome Fund.

## Data Availability

The data underlying this article are available in the article and in its online supplementary material.

## Author Contributions

LS performed all experiments and analyses, except where otherwise noted, and drafted and edited the manuscript. KTD assisted with topology tests. JFW set up the HMMER pipeline and assisted with interpretation. SDK performed enterobactin detection assays. CG and MG provided reagents and advised on their use. DAO, ALL, XZ, and XXS provided genome sequences and annotations. LS, AR, and CTH designed and conceived the study. AR and CTH secured funding and edited the manuscript. All authors approved the manuscript.

---

**Figure S1.**
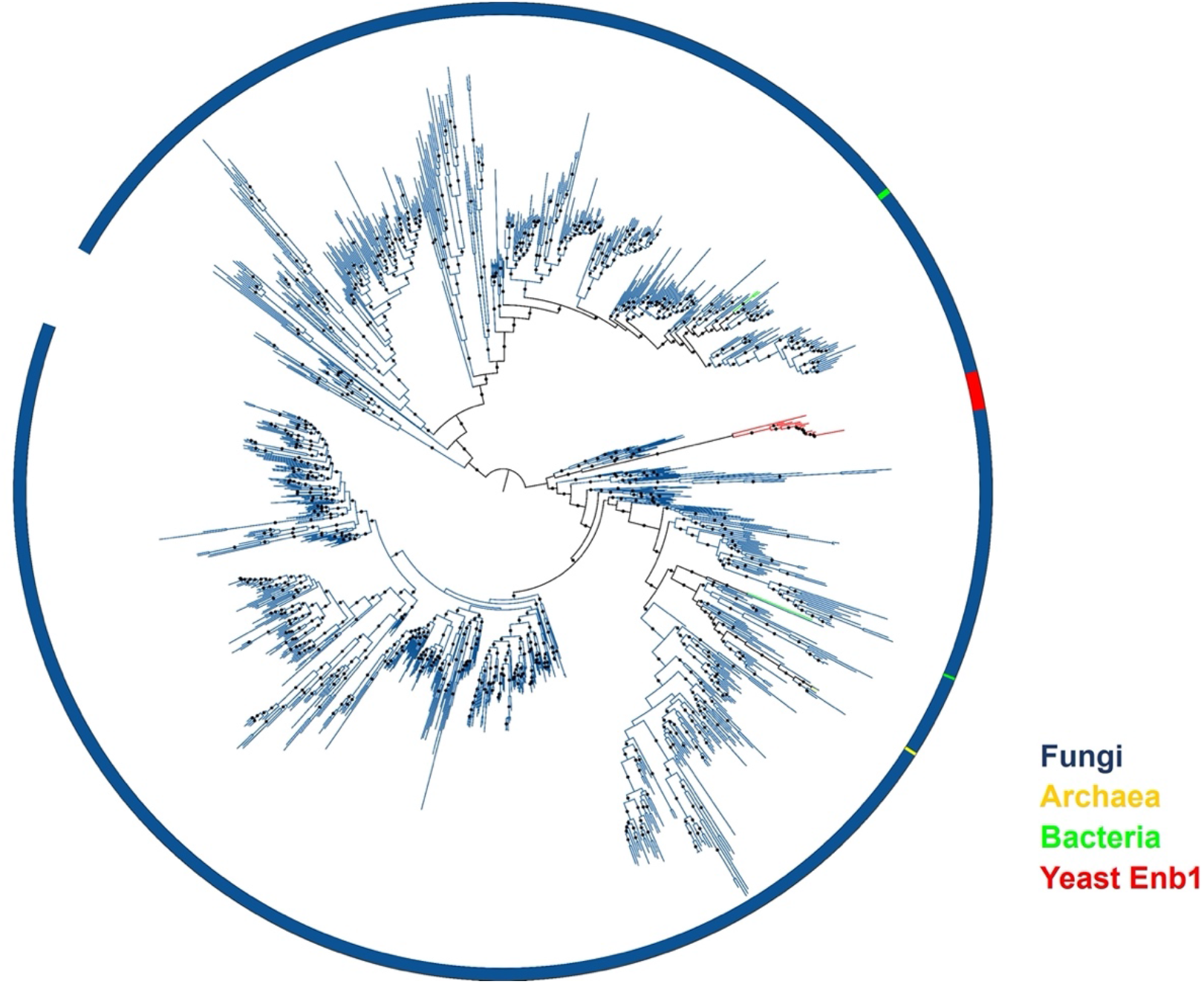
Midpoint-rooted maximum likelihood phylogeny of top Enb1 BLASTp hits from NCBI nr database. The maximum likelihood (ML) phylogeny was constructed with 1,165 sequences obtained from the 2,626 top BLASTp hits with *E*-values < e-60 by removing sequences with > 90% similarity using the clustering function of the mmseq2 (Steinegger and Söding 2017). The branches and outer strip were colored based on the key on the bottom right. Nodes with bootstrap values > 90% were highlighted using (•).

**Figure S2.**
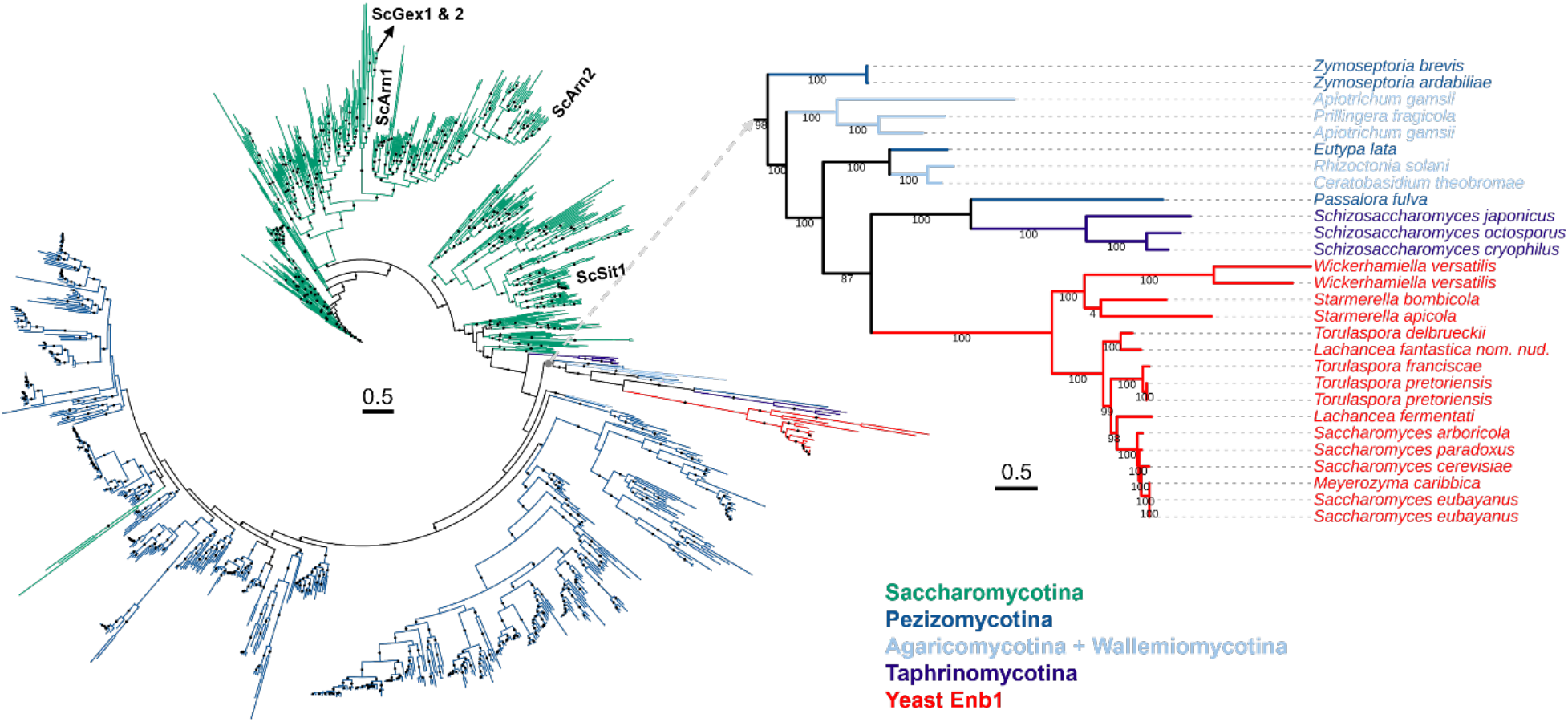
Midpoint-rooted maximum likelihood amino acid phylogeny of Enb1 homologs, including paralogs, from a database of 1644 fungal genomes. The branches were colored based on the key on the bottom right. Nodes with bootstrap values > 90% were highlighted using (•). Other Arn family proteins from *S. cerevisiae* and *St. bombicola* are labeled. The monophyletic clade consisting of Enb1 proteins from yeasts is zoomed in with the pruned tree.

**Figure S3.**
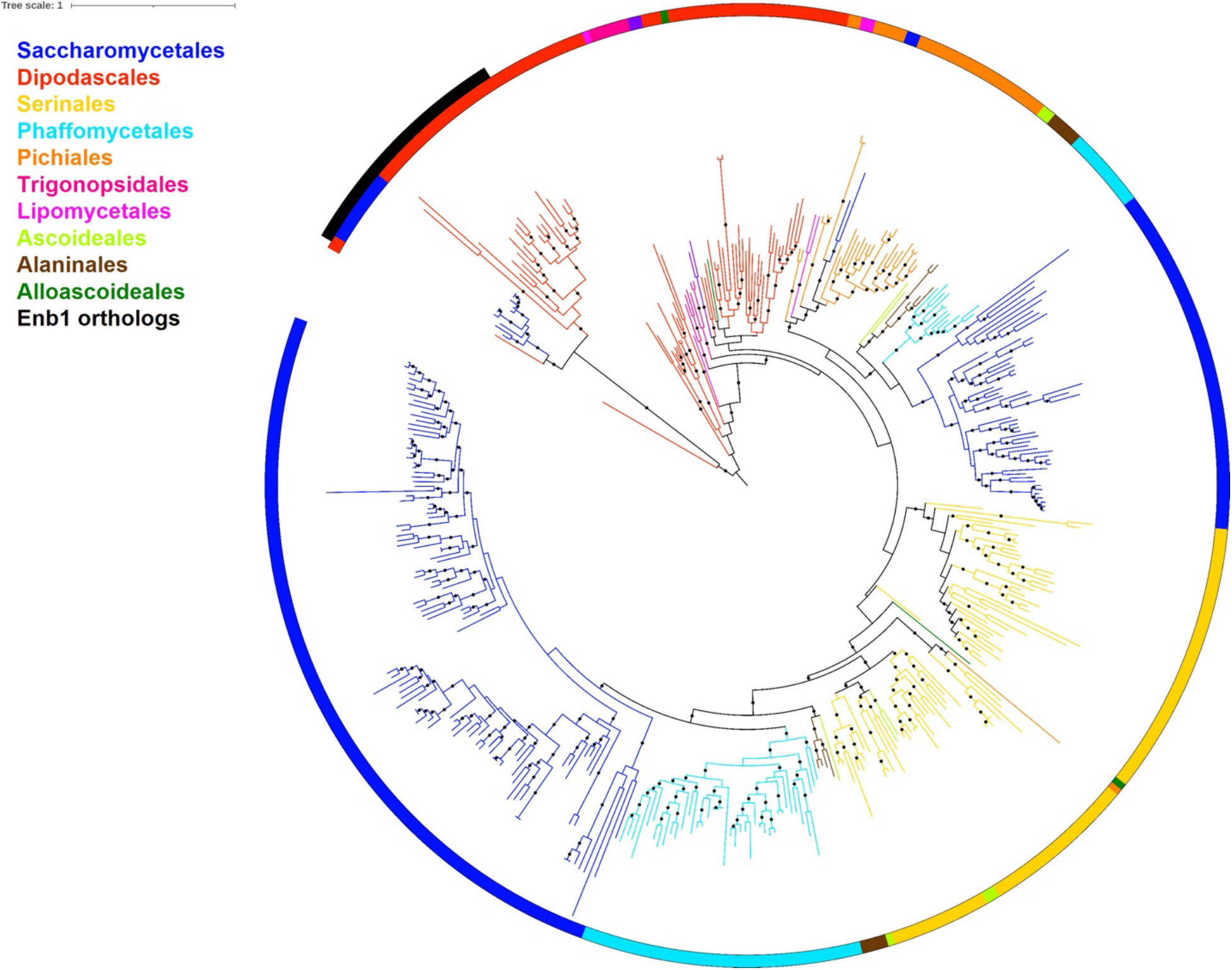
Midpoint-rooted maximum likelihood phylogeny of Enb1 homologs, including paralogs, from the yeast subphylum Saccharomycotina using an amino acid sequence alignment. Enb1 orthologs are marked in black. Nodes with bootstrap values > 90% out of 1000 bootstrap replicates were highlighted using (•). Yeast major clades are colored according to (Shen et al. 2018) and are now circumscribed as orders (Groenewald et al. 2023).

**Figure S4.**
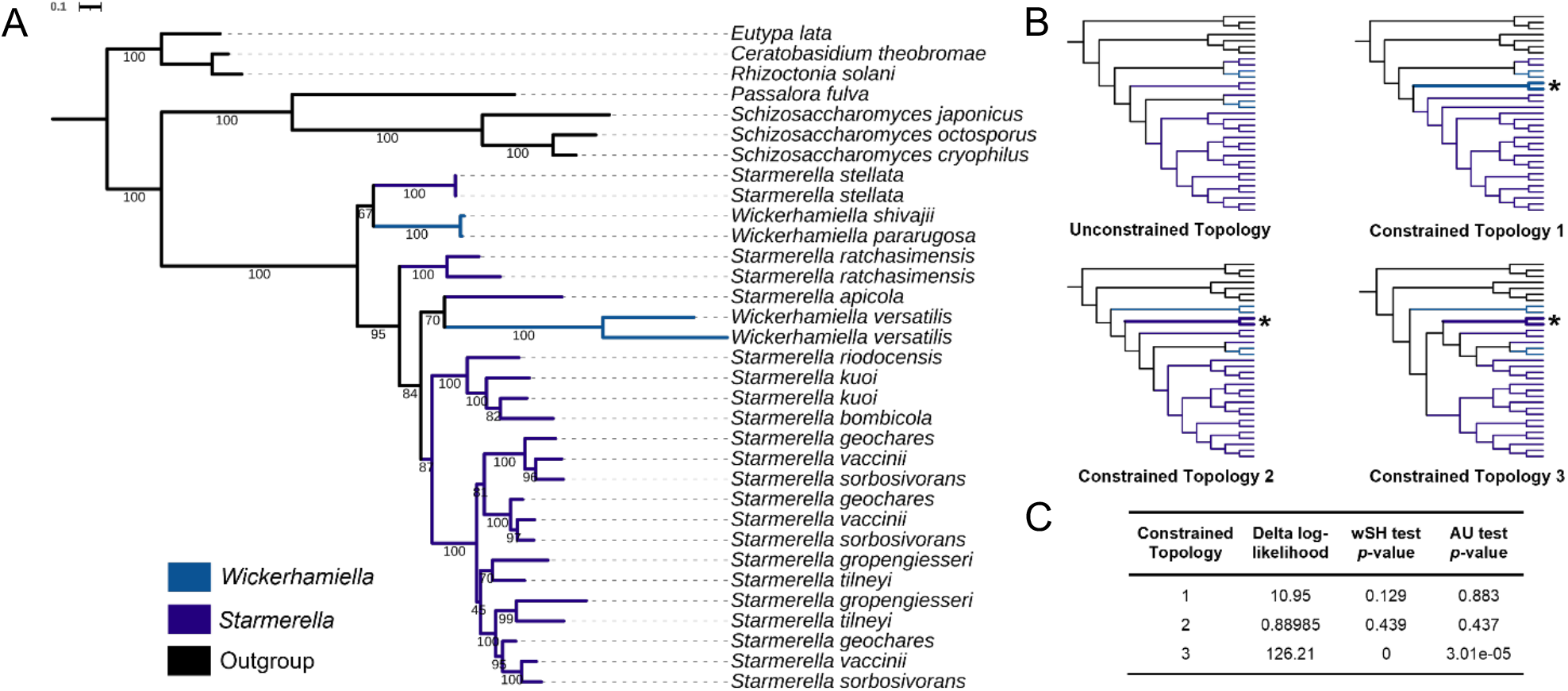
Topology tests of possible HGTs within the W/S clade yeasts. Maximum likelihood phylogeny of all W/S clade Enb1 protein sequences (A). The 7 closest relatives of Enb1 proteins from filamentous fungi were chosen as outgroups. Branch values shown are the percentages of branch support out of 1000 bootstrap replicates. Branches are colored in light blue, dark blue, and black to represent species from the *Wickerhamiella*, *Starmerella*, and outgroup lineages, respectively. Branches with the same species name occur because some genomes encode multiple copies of Enb1. The simplified topologies of the unconstrained and constrained gene trees are shown in (B). Branches repositioned in the constrained topology are denoted with (*). Constrained Topology 1 enforced the Enb1 homologs of *W. versatilis* to be outgroups to the Enb1 homologs of all the *Starmerella* species, except for *St. stellata*. Constrained Topology 2 enforced the Enb1 homologs of *St. stellata* to be outgroups to the Enb1 homologs of all other *Starmerella* species. Constrained Topology 3 enforced the clustering of the Enb1 homologs from *St. stellata* with those from the *St. ratchasimensis*, *St. apicola*, *St. riodocensis*, *St. kuoi*, and *St. starmerella* to conform to the species phylogeny. The delta log-likelihoods of constrained topologies are compared to the unconstrained topology, and the SH and AU topology test results are shown in (C).

## Supplementary Datasets

**Dataset S1-1.** Top hits of BLASTp against the NCBI non-redundant protein sequences database using *St. bombicola* Enb1 protein sequence as query.

**Dataset S1-2.** Top hits of BLASTp against the NCBI non-redundant protein sequences database, excluding fungi, using *St. bombicola* Enb1 protein sequence as query.

**Dataset S2.** Top hits of HMMER scan against the 1644-fungi genome database using the aligned sequences of *S. cerevisiae* and *St. bombicola* Enb1 proteins as queries.

**Dataset S3.** Top hits of HMMER scan against the 345-yeast genome database using the aligned sequences of *S. cerevisiae* and *St. bombicola* Enb1 proteins as queries.

**Dataset S4.** Newick format strings of phylogenetic trees constructed in this work.

**Dataset S5.** Strains used in this study.

**Dataset S6.** Oligonucleotides for construction of *St. bombicola* deletion mutants.

